# Disulfide bridge formation prevents CaMKII/Calmodulin interaction in Parkinson’s disease

**DOI:** 10.1101/2020.02.14.947960

**Authors:** Roberto Di Maio, Ignacio J. General, Emily Furbee, Joseph C. Ayoob, Sandra L. Castro, Ivet Bahar, J. Timothy Greenamyre, Filippo Pullara

## Abstract

There is increasing evidence for disordered Ca^2+^ signaling in dopamine neurons in Parkinson’s disease (PD), and this likely involves altered Ca^2+^/calmodulin-dependent protein kinase II (CaMKII) function. Previous work suggests that oxidative stress - a major feature in PD pathogenesis - affects regulatory methionine residues that sustain CaMKII activity in a Ca^2+^/CaM-independent manner. Here, applying computational modeling, we predicted formation of a defined disulfide bridge close to the CaMKII docking site for Ca^2+^/CaM binding. *In vitro* and *in vivo* investigations using PD models revealed formation of a disulfide bridge and loss of the CaMKII–calmodulin interaction. Mutagenesis of the relevant cysteine residues abrogated disulfide bridge formation and recovered the CaMKII–calmodulin interaction. Importantly, dopamine neurons from post-mortem PD brain specimens also lost this regulatory protein-protein interaction, providing relevance in the human disease. This study provides novel insights into oxidative CaMKII-CaM dysfunction, which may contribute to the pathophysiology of PD.

---

Ca^2+^/calmodulin dependent protein kinase II, CaMKII, is a serine/threonine specific kinase regulated by the Ca^2+^/calmodulin (Ca^2+^/CaM) complex. CaMKII is involved in a large number of biological processes, including Ca^2+^ regulation [1], the cell cycle [2], and T-cell activation [3]. CaMKII also regulates neuronal processes, such as long-term potentiation (LTP) and the modulation of neuronal excitability.

There are 4 CaMKII genes in humans, α, β, γ and δ, which have high sequence identity and are expressed in at least 38 isoforms [4] through alternative splicing. In the brain, CaMKIIα performs its function as a dodecameric protein holocomplex. This complex is composed of two stacked rings of six subunits each. Each subunit of the system is composed of three subdomains (Figure 1): 1. the C-terminal hub, located towards the center of the ring; 2. the N-terminal kinase head; and, 3. a linker that connects the hub with the kinase head. In the dodecameric complex, when the hub and kinase head are in close proximity, the protein conformation is called “closed” or “auto-inhibited,” because it is not bound to the Ca^2+^/CaM complex. Conversely, when the hub and kinase head are far apart, the linker is extended in the active “open” conformation, and CaMKIIα may bind the Ca^2+^/CaM. The mechanism governing the transition between the autoinhibited and active conformations is modulated by the phosphorylation of several amino acids. However, the molecular mechanisms of this regulation are still not well understood.

**Figure 1.**
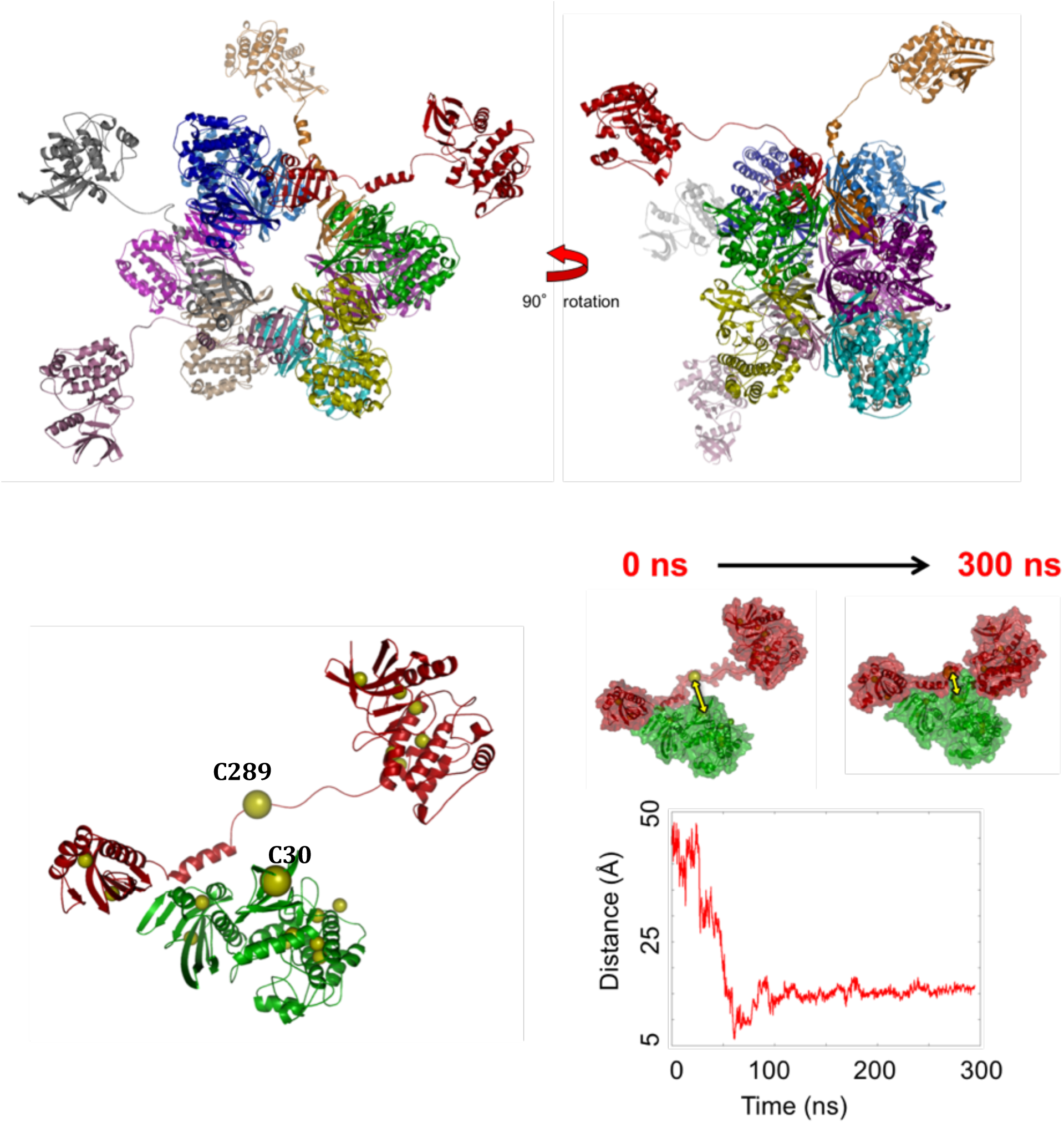
**A**: Top and side views of the full dodecamer model of CaMKIIα. In this conformer there are four CaMKII subunits in the open conformation and the remaining eight in the closed conformation. **B**: A detailed view of two adjacent CaMKII subunits. Represented as yellow spheres are the Cα of the cysteines. The Cα of C30 and C289 are represented by two larger yellow spheres. **C:** Snapshots from one of the all atoms molecular dynamics simulations highlighting the open and closed conformations. The double yellow arrows indicate the Cα-Cα distance between C30 and C289. The graph shows the time evolution of the distance between those two residues. Note that the minimum Cα-Cα distance reached during this simulation is less than 6Å.

Other conformational changes in CaMKIIα or modifications such as disulfide (S-S) bridge formation may play a role in this regulation. The latter is particularly intriguing given that both CaMKIIα dysfunction and oxidative stress, which can induce S-S bridges in proteins and protein complexes, are both implicated in neurodegenerative diseases. However, despite the large amount of information on both the physiopathology of CamKIIα in the brain and the ubiquitous presence of oxidative conditions accompanying neurodegeneration, the link between these two phenomena remains under-explored. Furthermore, there is no strong evidence of redox modulation of CaMKII activity/function in brain, though recent studies report redox sensitive properties of CaMKII in the heart and lung [5]. One reason for this is that currently there are no available experimental structures of CaMKIIα protein dodecameric complexes with subunits in the open and active conformation. The only structure available of CaMKII in an open conformation is a crystal structure of a CaMKIIδ monomer (PDB-ID 2WEL). The most complete CaMKII structure appeared in 2011 [6] and describes the structure of a CaMKIIα splice variant with all subunits in the closed/inactive conformation. A more complete structural analysis of CaMKII in various conformers and under different conditions, particularly redox modulation, would be an important next step in providing a more detailed mechanism of its regulation and deciphering the role of CaMKII in neurodegenerative diseases.

Dysregulation of Ca2+ homeostasis and signaling, and redox imbalance, have been suggested as key factors in various forms of neurodegeneration, including PD [7]. While most neuronal types use Na^+^ to generate action potentials, nigrostriatal dopamine (DA) neuronal activity is characterized by spontaneous firing that relies on Ca^2+^ currents mediated by CaV1.3 channels. In at least some neuronal populations, CaV1.3 forms a complex with CaMKII that modulates channel function [8].

The Ca^2+^-specific physiology of substantia nigra dopamine neurons requires precise regulation of certain systems, including CAMKII-CaM-mediated signaling and mitochondrial Ca^2+^ uptake to avoid elevated and harmful concentrations of cytosolic Ca^2+^. Such Ca^2+^ cycling and signaling is associated with ROS production. In addition, excess ROS may derive directly from DA metabolism, which generates harmful by-products, such as semiquinones and hydrogen peroxide [9]. The overall result is an increase in the basal levels of ROS in nigrostriatal neurons that reduces their ability to tolerate further oxidative insults. In this context, systemic administration of rotenone or paraquat, which causes ROS production, and which targets every cell in central nervous system, results in selective damage of the DA system, and, reproduces aspects of PD pathogenesis [10, 11].

In this study, we undertook a computational modeling approach to probe the dynamics and regulation of a CaMKII dodecameric complex. Here we provide evidence of oxidatively-induced formation of a functional S-S bridge in the docking site of CaMKII for Ca^2+^/CaM complex, which interferes with the normal CaMKII-Ca^2+^/CaM interaction. Computational approaches and mutagenesis experiments confirmed that residues C30 and C289, belonging to two distinct and adjacent CaMKII subunits, form this bridge. Moreover, we provide evidence of an oxidatively-mediated, drastic reduction of the CaMKII-Ca^2+^/CaM interaction in human idiopathic PD and in the rotenone rat model thereof, suggesting that this phenomenon may be relevant in PD pathogenesis.

## Results

Examining the structural dynamics of CaMKIIα required a structure with protein subunits in the open/active conformation within the dodecameric complex. In the absence of such a structure, we used existing crystal structures of CaMKIIα in the closed conformation and CaMKIIδ in an open configuration to create a model of a CaMKIIα dodecamer with four open subunits and eight closed subunits (see Methods). We ran 8 independent all atoms Molecular Dynamics simulations with explicit water (see Methods) of the modeled dodecamer, for a total simulation time of around 2.6μs. We evaluated the distance between the α-carbons of all of the cysteines and identified two cysteines, C30 and C289, belonging to adjacent subunits that get very close to each other due to the overall motions of the protein complex (Figure 1 B-C).

The analysis of the Cα-Cα distances of C30 and C289 in independent simulations shows different outcomes emphasizing the stochasticity of the phenomenon. We found that in 4 cases the close proximity between Cαs (< 10Å) is transient and it survives for just few ns. In two of the simulations we ran, the C30-C289 interaction is rare or does not occur. Finally, in the remaining 2 cases, the distance between those two cysteines stabilizes around 10-15Å for a significant period of time (Figure 1 B-C).

In most of the simulations, after around 100ns, the Cα-Cα distance drops to less than 20Å with a minimum separation below 6Å. We observe in our modeling that C30-C289 remain generally in close proximity. This suggests that in oxidative stress conditions, which is a hallmark of neurodegenerative diseases, it is possible that the interaction between the thiol groups of C30 and C289 will lead to a S-S bridge formation. Moreover, when C30 and C289 are in close proximity, they occlude the region where CaM binds (Threonine residues 305 and 306) [12, 13], which is otherwise accessible (Figure 2). Additionally, the Threonine residue, T286, whose phosphorylation is known to be one of the first steps in CaMKII activation [14-16], would similarly be strongly perturbed if a S-S bridge were to be formed between C30 and C289. These computational results suggest the presence of a S-S bridge between adjacent CaMKII subunits in the CaM biding region, which apparently would disrupt CaMKII-Ca^2+^/CaM binding, as well as CaMKII function.

**Figure 2.**
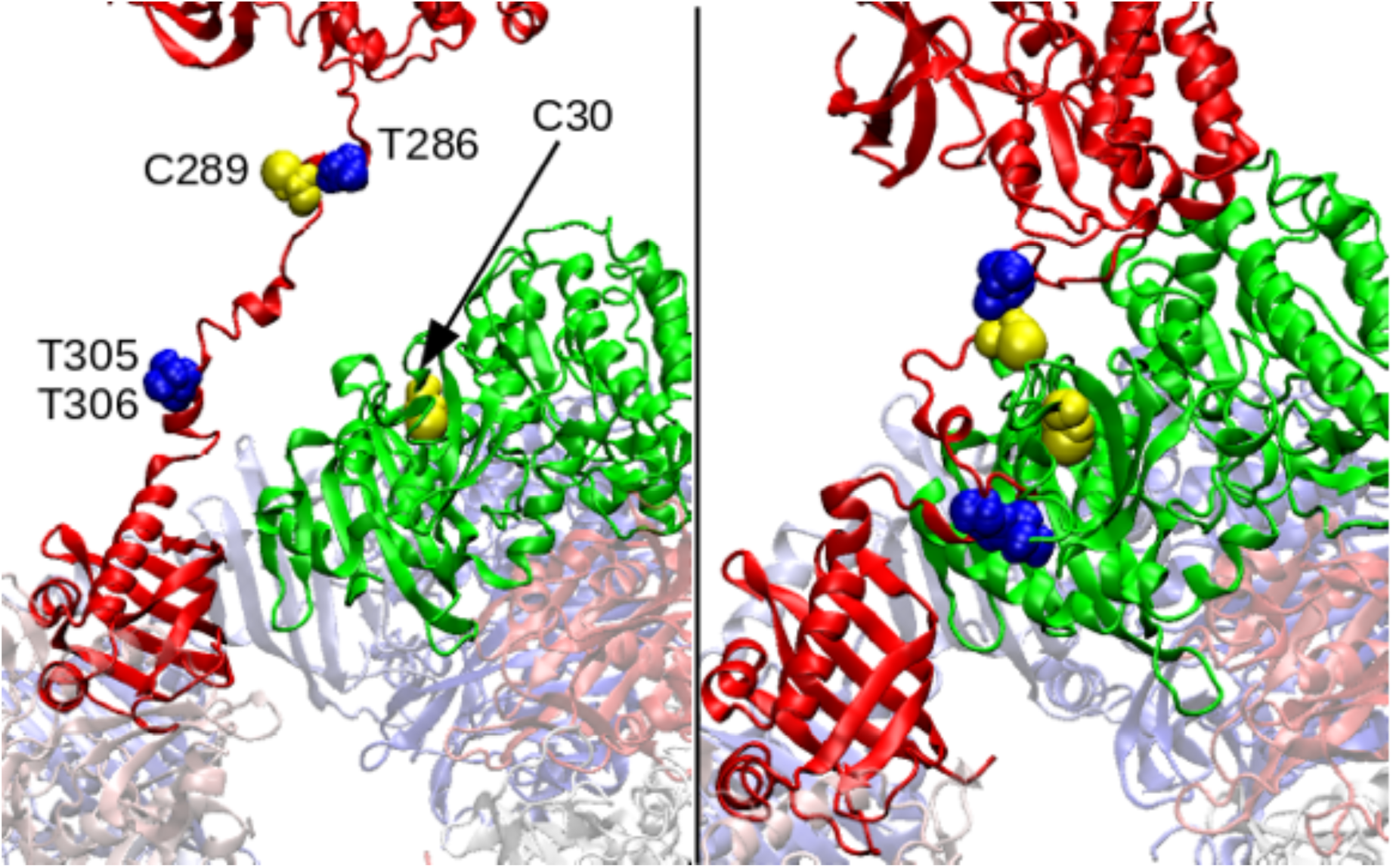
Close-up view of two subunits of CaMKII, showing the location of C30 and C289 (S-S bridge), T305 and T306 (binding region of CaM), and T286 (involved in CaMKII activation). When the bridge is not formed (left panel), there is room around T305/306 for CaM to bind. When the bridge is formed (right panel), the accessible area to the binding site gets greatly reduced, possibly preventing CaM binding.

To support our computational findings of a putative S-S bridge between adjacent subunits of CaMKII, we examined whether such a bond is formed under oxidative conditions in ventral midbrain primary neurons. We performed “redox western blot” assays, which allow detection of thiols derived from S-S bridges via an alkylating reaction with a 10 kDa PEG-NEM tag, resulting in an observable 20 kDa shift of the predicted molecular weight for each S-S bridge. In the absence of an oxidizing treatment, no significant shift of CaMKII is observed, whereas in cells exposed to the mitochondrial complex I inhibitor, rotenone, a 20 kDa shift was observed for CaMKII (Figure 3 A-B), suggesting the formation of a single S-S bridge in CaMKII under oxidative stress conditions.

**Figure 3.**
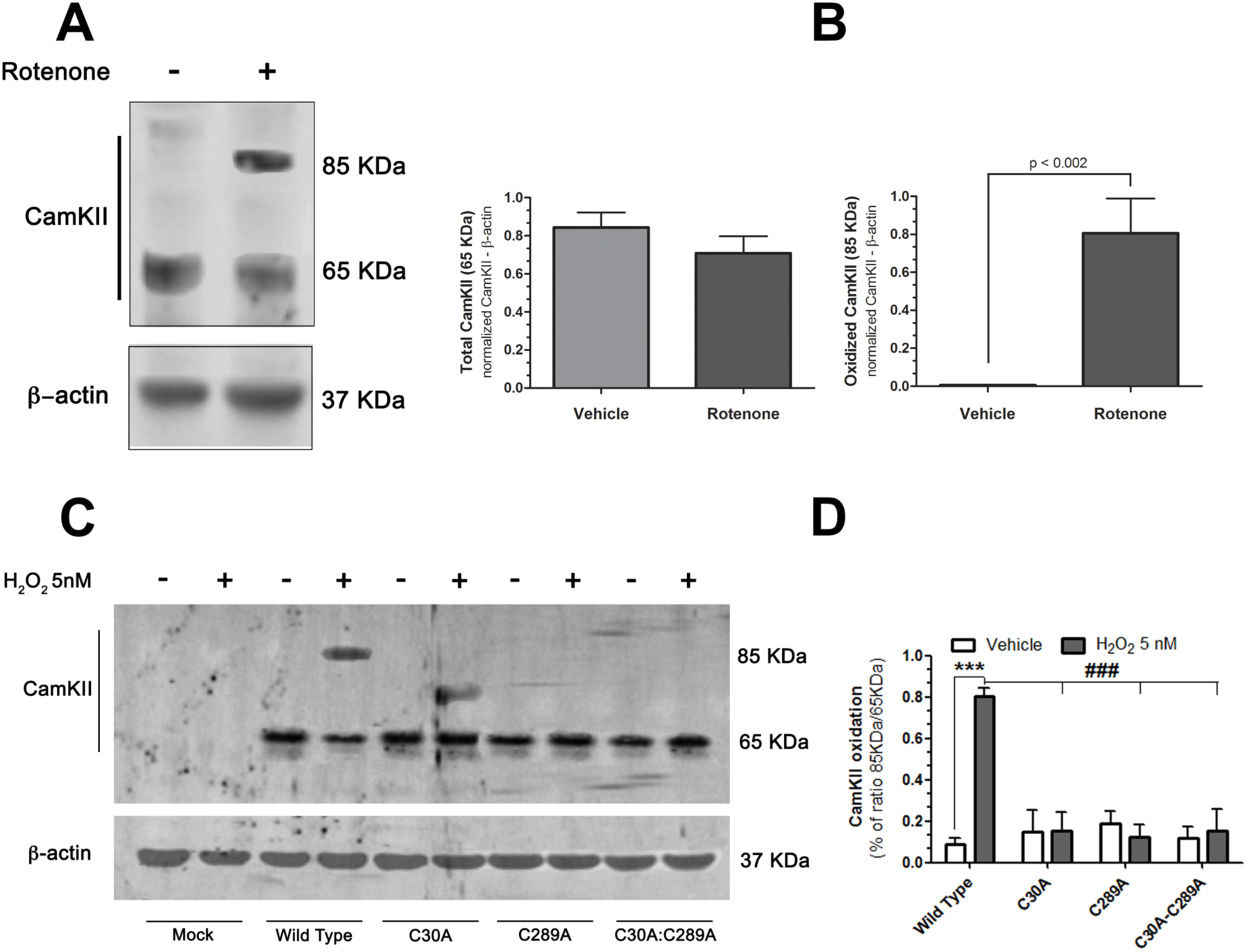
Oxidative stress-related disulfide formation in CaMKII. **A**: Redox WB for CamKII revealed a 20KDa MW shift in lysates of ventral midbrain cultures exposed to 50nM rotenone for 24hours, suggesting the formation of a disulfide bridge. **B**: Quantification of the intensity of the bands shows no significative changes in CamKII expression and a significant increase of CaMKII in oxidized state (85KDa band) in H_2_O_2_ treated cells (2-tailed unpaired t-test with Welch’s correction; p < 0.002; n=3). **C**: Consistent with the results obtained in ventral midbrain cultures, redox WB of lysates from *Drosophila* cells expressing human wild type CamKIIα showed the formation of a disulfide bridge when cells were exposed to 5nM of H_2_O_2_ for 24 hours. The phenomenon was prevented in CamKII C30A and C289A mutants. Note that H_2_O_2_ treatment of cells expressing only the mutant C30A elicited the appearance of a 10 KDa shift of CaMKIIα, suggesting an oxidative process exerted only on C289. This may be explained by the computational evidence that C289 is more exposed to the solvent (see Figure 1, bottom panel: C289 is in the linker of the red subunit). Therefore, although C289 cannot form a S-S bridge with C30 in C30A mutant, its high exposure facilitates other kinds of thiol oxidation, such as S-NO formation. **D**: Quantification of band intensity expressed as ratio 85/65 KDa (ONE-WAY ANOVA + Bonferroni test; p < 0.0001; n=3)

To identify which cysteine residues are involved in this S-S bridge, we set out to directly test our computational results that implicate C30 and C289 as the bridge-forming pair. We performed mutagenesis on human CaMKIIα and expressed these mutant transgenes in a *Drosophila* S2R+ adherent cell line, to avoid cross-immunoreactivity with native proteins. Redox western blot assay in S2R+ wild type, C30A, C289A, and C30A:C289A mutants revealed that exposure to 5 nM H_2_O_2_ elicited formation of an S-S bridge (20 kDa MW shift in wild type CaMKIIα), similar to that seen in the ventral midbrain neurons (Figure 3C-D). There was no MW shift observed under basal conditions. Likewise, no MW shifts were observed with either mutant alone, or in combination, under normal or oxidative stress conditions, adding strong support for a specific C30/C289-related S-S bridge in CaMKIIα.

In addition to revealing an S-S bridge in CaMKIIα, our computational studies also indicate that Ca^2+^CaM binding to CaMKIIα is likely to be blocked when the S-S bridge is formed between C30 and C289. To test this, we employed a proximity ligation assay, which provides fluorescent signal where there is interaction between two tagged proteins, in this case CaMKIIα and CaM. In rat ventral midbrain neurons, under basal conditions, we observed a strong CaMKIIα:CaM interaction (Figure 4 A-B). Under oxidative stress conditions induced by rotenone, the interaction between CaMKIIα and CaM was significantly reduced. We repeated these experiments in the *Drosophila* S2R+ cell line using C30A and C289A mutations, alone and in combination, to test the potential role of these cysteine residues in the CaMKIIα:CaM interaction. In both normal and oxidative stress conditions, the CaMKIIα:CaM interaction was only disrupted when the wild type protein was expressed in H_2_O_2_-treated cells (Figure 4 bottom-D). These results demonstrate that normal CaM binding to CaMKIIα is disrupted when the S-S bridge is formed between C30 and C289 when cells are under oxidative stress.

**Figure 4.**
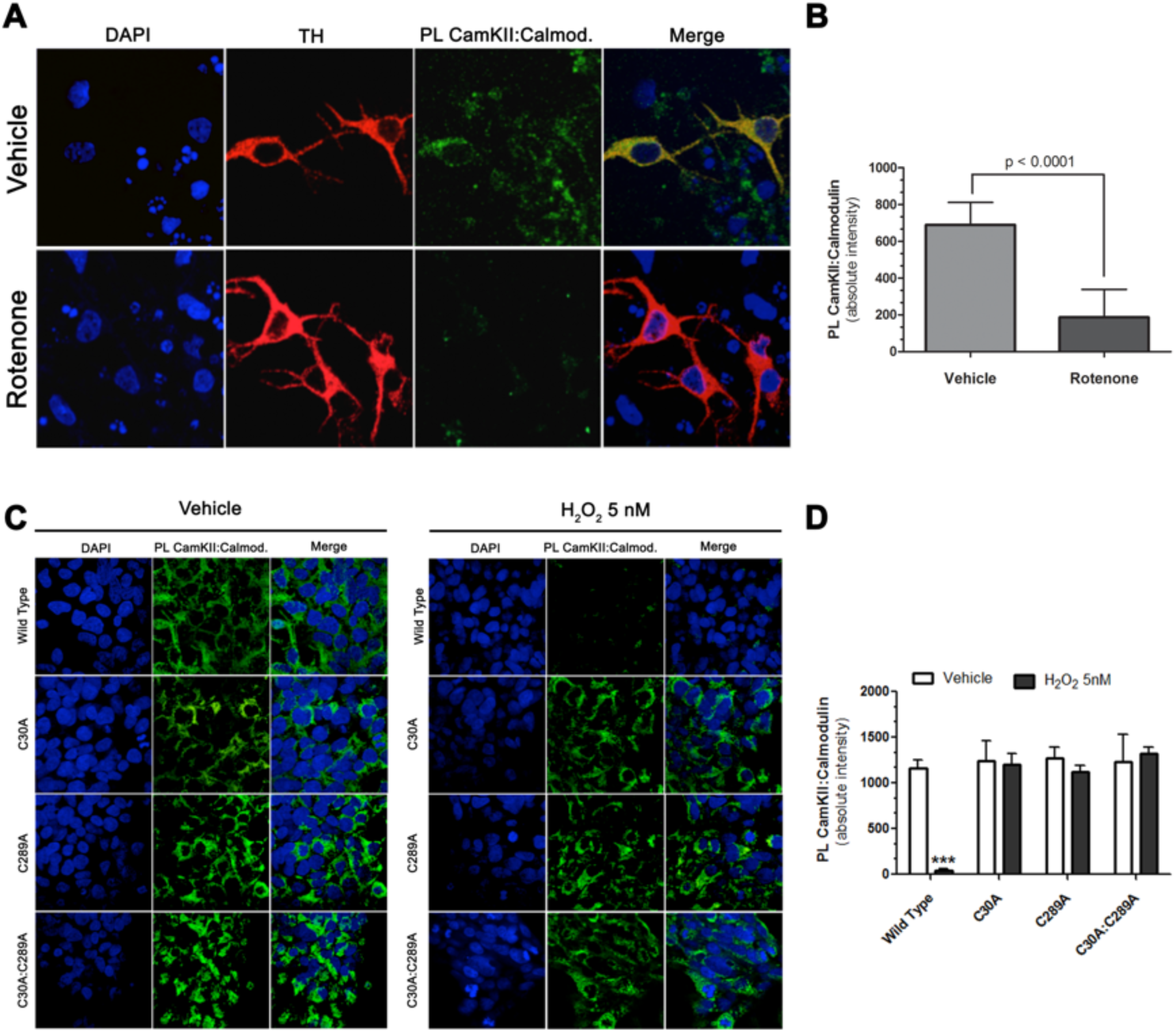
Disulfide formation between C#) and C289 prevents CaMKII/CaM interaction. **A:** Proximity ligation between CamKII and calmodulin in DA neurons of ventral midbrain primary cultures. In vehicle-treated cells, there is intense punctate signal for CamKII and calmodulin interaction. In contrast, rotenone treatment induced loss of fluorescence PL signal for CaMKII-Ca^2+^/CaM **B:** The plot reports the mean cellular PL fluorescence values (vehicle- or rotenone-treated). In each sample, PL signal was measured in 20-40 TH-positive neurons. Statistical testing by 2-tailed unpaired t-test with Welch’s correction. **C:** Drosophila cells expressing human CamKII wild type, C30A, C289A or the double mutation C30A-C289A were exposed to 5nM H_2_O_2_ for 24 hours. PL assay for CaMKII-Ca^2+^/CaM revealed, as in VMB cultures, the loss of signal for CaMKII-Ca^2+^/CaM interaction. Noteworthy, all the mutations were able to prevent H_2_O_2_-mediated CaMKII-Ca^2+^/CaM disruption, suggesting the critical role of the redox status of C30 and C289 in modulating the functional binding of Ca^2+^/CaM. **D:** Quantification of cellular PL fluorescence CaMKII-Ca^2+^/CaM For each sample, PL signal was measured in 50–70 cells and the data are from 3 independent experiments. Statistical testing by ONE-WAY ANOVA with Bonferroni post-hoc test.

The mitochondrial complex I inhibitor, rotenone, reproduces many features of PD-related pathogenic events including mitochondrial dysfunction, oxidative stress, altered Ca2+ homeostasis, and accumulation of toxic forms of α-synuclein [17-20]. We further assayed the CamKIIα:Ca^2+^CaM interaction in this paradigm to examine if the disruption of binding is manifested in the *in vivo* rat PD model. A robust CamKIIα:Ca^2+^CaM interaction was observed in nigrostriatal neurons under basal conditions, but, we observed a significant reduction of CamKIIα:Ca^2+^CaM interaction (by over 70%) in dopamine neurons of substantia nigra pars compacta (SNpc) from rotenone-treated animals, which were exhibiting PD-like behaviors (Figure 5). Our results from rotenone-treated rats demonstrated an impaired CaMKII/CaM interaction. Moreover, previous evidence in PD models report CaMKII hyperactivity related to high levels of phosphorylation at the Thr 286 residue in in parallel with decreased activity of the specific phosphatase PP1_ϒ_1 [21, 22]. The event may be linked to an uncontrolled Calmodulin-independent CaMKII activity. Similarly, as (QQQ Add supplementary FIGUREfig S…) shows, we detected a significant increase of pThr286 in rotenone treated animals, suggesting that, in absence of CaM modulation, CaMKII exerts an uncontrolled Kinase activity. To determine whether this process is relevant to idiopathic human PD, we performed PL assays (CaMKII:CaM) in blinded postmortem substantia nigra sections from PD patients (N=5) and from controls (N=4). Compared to controls, nigrostriatal dopamine neurons from all PD cases showed very low PL signal, indicating loss of interaction between CaMKII and CaM (Figure 6). This evidence strongly suggests that CaM-mediated modulation of CaMKII is impaired in the human disease and may be a relevant factor in PD pathophysiology. Together, these results provide a strong case for an oxidative stress-induced S-S bridge in CaMKII that occludes calmodulin binding and contributes to the pathogenesis of PD.

**Figure 5.**
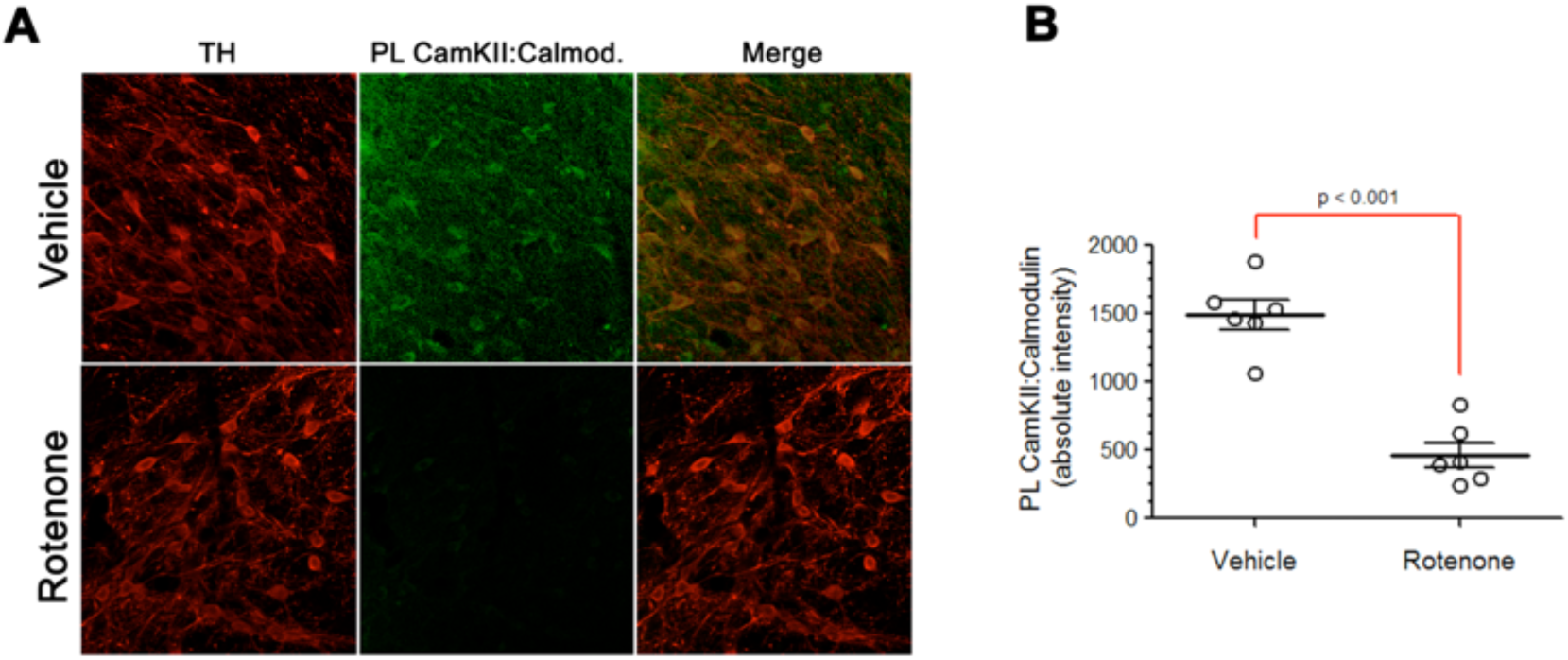
Evidence of disrupted CaMKII/CaM interaction in PD model of rotenone. **A:** In a vehicle-treated rat (top row), a strong PL signal for CaMKII-Ca^2+^/CaM interaction was detected. In contrast, rotenone treatment (bottom row), prevented PL signal CaMKII-Ca^2+^/CaM, suggesting, again that, under mitochondrial injury and oxidative stress conditions, the two proteins lose the ability to interact. **B:** Quantification of PL fluorescence values for individual animals (vehicle- or rotenone-treated). Each circle represents the mean of PL fluorescence for a single animal. In each animal, PL signal was measured in 35–50 nigrostriatal neurons per hemisphere. Statistical testing by 2-tailed unpaired t-test with Welch’s correction.

**Figure 6.**
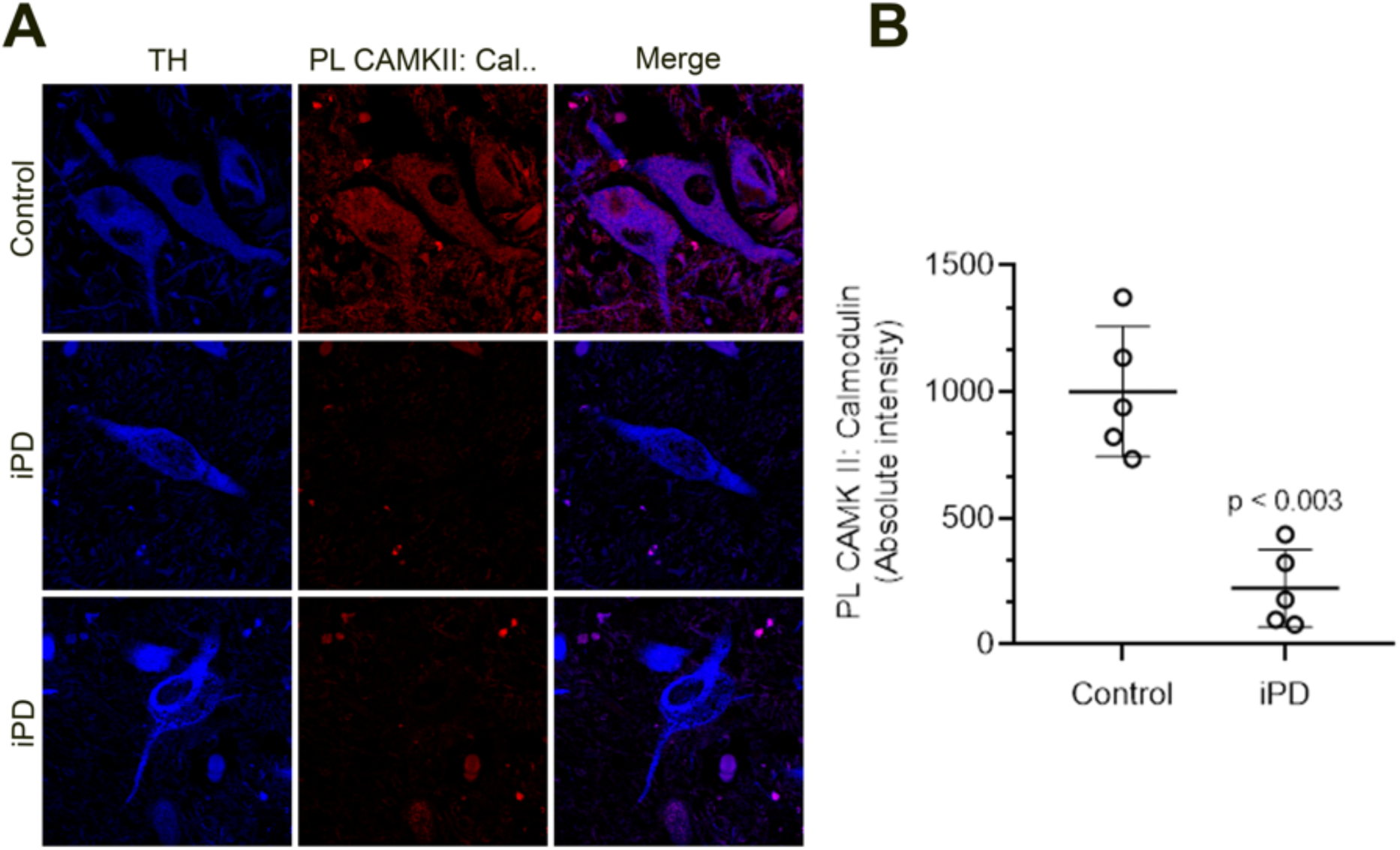
Relevance of CaMKII/CaM interaction in idiopathic PD Proximity ligation between CamKII and calmodulin in human brain. **A:** In control patients (top row), a strong PL signal for CaMKII-Ca^2+^/CaM interaction was detected. In contrast, idiopathic Parkinson’s disease (iPD, bottom row), showed little PL signal CaMKII-Ca^2+^/CaM, suggesting, again that, under mitochondrial injury and oxidative stress conditions, the two proteins lose the ability to interact. **B:** Quantification of PL fluorescence values for individual patients (control or iPD). Each circle represents the mean of PL fluorescence for a single patient. In each patient, PL signal was measured in 35–50 nigrostriatal neurons. Statistical testing by 2-tailed unpaired t-test with Welch’s correction.

## Discussion

In this study, we applied novel computational techniques in conjunction with experimental *in vitro* and *in vivo* approaches to provide new insights into the mechanisms of regulation of CaMKIIα and its interaction with calmodulin. Our computational results identified the presence of a putative S-S bridge across two adjacent CaMKIIα subunits in its dodecameric configuration. Interestingly, this predicted bridged conformation occludes the binding site for calmodulin as well as the activation site for CaMKIIα. In cell culture models, including ventral midbrain primary cultures, we confirmed the presence of this S-S bridge, which only occurs under oxidative stress conditions, and identified the specific cysteines involved. Further investigation demonstrated that formation of this bridge does in fact block CaMKII□ interactions with calmodulin, and again this only occurs in oxidative stress conditions. Mutating one or both cysteine residues that form the bridge both blocks S-S formation and restores calmodulin binding. In this context, we observed a significant loss of interaction CaMKIIα/calmodulin in *substantia nigra* dopamine neurons of rotenone-treated animals and, most importantly, in dopamine neurons in post-mortem PD brain specimens. Our results, obtained with complementary experimental approaches, demonstrated for the first time a specifically-defined S-S bridge in CaMKIIα that affects proper modulatory control of CaMKII exerted by Ca^2+^/calmodulin complex. This novel mechanism of CaMKIIα regulation under oxidative stress conditions, provides insights into the potential relevance of CaMKIIα dysfunction that may trigger a downstream pathological cascade of events culminating in PD-related neurodegeneration. Further studies are required to better define the role of oxidative CaMKII dysfunction in the pathophysiology of PD.

## Methods

### Molecular Dynamics Computer Simulation

Computational simulations of the CaMKIIα were performed using the AMBER15 [23] software package, with all of them—except for the minimizations—using the GPU version of the PMEMD program. The crystal structure of CaMKIIα in a closed/auto-inhibited conformation (PDB structure 3SOA) was used as the starting structure [6] of human CaMKII in its α isoform, with a β7 linker (the shortest linker in all known splice variants [24]. Some of the subunits in this dodecamer were converted computationally to open conformations based on the 2WEL PDB crystal structure of the CaMKIIδ isoform, which is in the open conformation. This was feasible since the kinase domains of both isoforms have a high sequence similarity of about 93%. The final structure contained eight closed and four open sub-units of CaMKIIα in a dodecameric configuration (Figure 1).

In order to bring the modeled molecule to a stable conformation, we ran several cycles of minimization and equilibration. We employed the Amber12 force field and the TIP3P water model. The systems were kept at a temperature of 298 K, using Langevin dynamics with a collision frequency of 2 ps^-1^, and part of the protocol used pressure control, via a weak-coupling Berendsen barostat, with a relaxation time of 2 ps. The SHAKE algorithm was adopted, allowing the use of a 2 fs time step. The protocol followed for minimization and equilibration was: 1) 100 cycles of minimization, using the XMIN method, followed by 5,000 cycles using steepest descent and another 5,000 steps using conjugate gradient; 2) 1 ns of heating to 298 K, followed by another ns at constant T and P (1 atm), and finally 20 ns with constant T. The simulation box consisted of one CaMKIIα dodecamer complex with eight subunits in the closed conformation and four subunits in the open conformation. All atom simulations were performed on CaMKIIα in a box with explicit water plus ions to neutralize electrostatic charges. Systems were >1.3×10^6^ atoms and all runs were >300ns. For all the runs we used Nvidia GeForce TITAN-X GPUs.

### Experimental

#### Ventral midbrain primary cultures

Neuronal cultures were prepared from embryonic day 17 Sprague-Dawley rats (Charles River, Wilmington, MA, USA). Embryos were obtained from two to three pregnant dams. Pooled ventral midbrain tissue was dissociated with enzymatic digestion using trypsin followed by mechanical trituration. Cells were seeded at a density of 5×10^5^/well in a 24-well plate. The cultures were maintained at 37 °C in a humidified atmosphere of 5% CO_2_ and 95% air in MEM with 2% FBS, 2% HS, 1 g/L glucose, 2 mM GlutaMax, 100 lM non-essential amino acids, 1 mM sodium pyruvate, 50 U/mL penicillin, and 50 lg/mL streptomycin. After 48 h, the culture medium was replaced with serum-free Neurobasal medium containing 0.5 mg/mL AlbuMAX I, 2 mM GlutaMAX I, 2% B27 supplement, 50 U/mL penicillin, and 50 lg/mL streptomycin. 50 ng/mL GDNF (#512-GF-050; R&D Systems. Minneapolis, MN, USA) was also added. Five days after seeding, the media was removed and replenished with Neurobasal medium. At 9 days in vitro (DIV), fresh Neurobasal medium was added over old medium in combination with GDNF. At 13 DIV, 70% of old medium was replaced. All the experiments were performed at DIV 15 exposing cells to Vehicle (DMSO) or 50nM rptenone for 24 hours. All procedures were performed with the approval of the University of Pittsburgh Animal Care and Use Committee.

#### Animals

Six-to seven-month-old adult male Lewis rats were used for all experiments (Envigo). The animals were maintained under standard conditions of 12 hours light/dark cycle in a 22 ± 1 °C temperature-controlled room with 50%-70% humidity. Water and food were provided ad libitum. Animals were adapted for two weeks to the described conditions before the experiments. All studies were approved by the Institutional Animal Care and Use Committee at the University of Pittsburgh and were performed in accordance with published National Institutes of Health guidelines.

#### Mutagenesis experiments

CaMKIIα and calmodulin expression in *Drosophila* cells was accomplished as follows: CaMKIIα and calmodulin coding sequences were copied from human cDNA clones CAMK2A transcript variant 2 (catalog SC109000, Origene, Rockville MD) and CALM1 Human calmodulin 1 transcript variant 1 (catalog SC115829, Origene, Rockville MD), using PCR primers to add flanking restriction sites, with the following primers: EcoRI-CAMK2A-FWD; GAGAGAATTCATGGCCACCATCACCTGCAC, CAMK2A-STP-SalI-REV; TCTCGTCGACTTAGTGGGGCAGGACGGAGGGCG, MfeI-CALM1-FWD; GAGACAATTGATGGCTGATCAGCTGACCGA, CALM1-STP-SalI-REV; TCTCGTCGACTCATTTTGCAGTCATCATCT. Products were ligated into appropriately gapped vector pRmHA3 (*Drosophila* Genomics Research Center, Catalog 1145) to generate pRmHA3-CaMKII and pRmHA3-CALM1. Point mutation variants (pRmHA3-CaMKII^C30^, pRmHA3-CaMKII^C289A^, and pRmHA3-CaMKII^C30A, C289A^) were generated by site directed mutagenesis of pRmHA3-CaMKII using QuickChange Lighting Kit (Agilent, Catalog 210518.) The full coding sequence of all pRmHA3-CaMKII constructs was verified by Sanger sequencing. pRmHA3-CaMKII constructs were introduced to *Drosophila* S2 cells (*Drosophila* Genomics Research Center, catalog 181) cultured under standard conditions in Schneider’s media with 10% FBS and Pen-Strep at densities between 0.5-2.0 × 10^6^, by transient Effectene transfection (Qiagen 301425) according to manufacturer recommendations. Expression was induced with 500 uM CuSO4 at the time of transfection and transfected cells were incubated for 48 hours under standard conditions. pRmHA3-mCherry was co-transfected with pRmHA3-CaMKII constructs to evaluate transfection efficiency and rule out morphological indications of toxicity in transfected cells. Oxidative stress was induced by treating S2 cells with 0.5 mM H_2_O_2_ in normal culture media for thirty minutes before lysis for immunoblots as described below.

#### *In vitro* and *in vivo* rotenone models of PD

Primary cultures were exposed to 50 nM rotenone (Sigma-Aldrich St. Louis, MO, USA) or vehicle (2% DMSO) for 24 hours. At the endpoint, cells were fixed in 4% paraformaldehyde. Cells were incubated overnight with primary antibodies for CaMKII, calmodulin, and tyrosine hydroxylase, as marker of dopaminergic neurons.

<u>For</u> *<u>in vivo</u>* <u>studies</u>, animals were randomly divided into vehicle (6 animals) and rotenone (11 animals) groups of treatment. Before injections, body weights were recorded for each animal. Rotenone was administered intraperitoneally once a day at a dose of 2.8 mg/kg until the end of the treatment (11-13 days). The solution was prepared as a 50X stock dissolved in pure dimethyl sulfoxide at final concentration of 2%, then diluted in Miglyol 812N (Sasol North America, Inc, Houston, TX; distributed by Warner Graham, Baltimore, MD, USA) at final concentration of 98%, and administered at 1 mL/kg. This regimen produces relatively uniform bilateral nigrostriatal lesions, leading to loss of about 50% of dopaminergic neurons [20]. Control animals received an equivalent amount of vehicle (2% dimethyl sulfoxide 98% Miglyol). Tissues were collected from each animal when they developed the debilitating behavioral phenotypes of akinesia, rigidity, and postural instability.

#### Histology

Animals were euthanized by CO_2_ inhalation followed by decapitation. The brains were removed following an intracardial perfusion with saline solution (NaCl 0.9%) and a fixing perfusion with 4% paraformaldehyde (PFA). After 48 hours, the brains were placed in 30% sucrose in phosphate buffered saline (PBS) for cryoprotection until infiltration was complete (at least 3 days).

#### Proximity Ligation Assay

Proximity Ligation Assay (PLA) (Duolink; Sigma Aldrich) was performed as described in https://reedd.people.uic.edu/ReedLabPLA.pdf in 4% PFA-fixed tissue or cell cultures to assess the level of interaction CaMKII-Ca^2+^/CaM under our experimental conditions. Samples were incubated with specific primary antibodies against CaMKIIα (Rb anti-CaMKIIα; ab92332 – abcam) and calmodulin (Ms anti-calmodulin; ab2860 – abcam). PLA probes consisting of secondary antibodies (anti-Rb and anti-Ms) conjugated with complementary oligos (plus and minus respectively) were added to the reaction and incubated. Ligation solution, consisting of two oligos and ligase, was added. In this assay, the oligos hybridize to the two PLA probes and join to a closed loop if they are in close proximity. An amplification solution, consisting of nucleotides and fluorescently labeled oligos, was added together with polymerase. The oligonucleotide arm of one of the PLA probes acts as a primer for “rolling-circle amplification” (RCA) using the ligated circle as a template, and this generates a concatemeric product. Fluorescently labeled oligonucleotides hybridize to the RCA product. The PL signal is visible as a distinct fluorescent spot. Fluorescent images were captured using a confocal microscope (Olympus BX61, Fluoview 1000; Melville, NY, USA) and quantification was carried out at 60-100X magnification. Control experiments included routine immunofluorescence staining of the proteins of interest under identical experimental conditions.

#### Redox Western Blot

Levels of oxidized thiols (S-S) in CaMKII were measured with a redox Western blot technique. Ventral midbrain primary cultures or Drosophila cells were lysed in 50mM Tris-HCl pH 7.0, 2% sodium dodecyl sulfate (SDS), 1mM EDTA, protease inhibitor cocktail and 100 µM N-ethyl-maleimide (NEM), to stably alkylate free thiol residues (SH). The lysate was heated at 65 °C for 5 min to denature proteins and incubated for 20 min at room temperature. Proteins were precipitated in ice-cold acetone to remove un-reacted reagents, resuspended in 50mM Tris-HCl pH 7, 2% SDS, and 10mM TCEP and heated at 65 °C for 5 min and incubated 20 min at room temperature to reduce S-S bridges to free SH groups. Proteins were precipitated again, and the pellet was re-suspended in 50mMTris-HCl pH 7, 2% SDS, and 100 µM polyethylene glycol-maleimide (PEG-NEM; 10KDa) to label the formerly oxidized thiols. After 20 min of incubation, the proteins were precipitated and a Western blot for CaMKIIα was performed. PEG-NEM labeling of formerly oxidized thiols increases the molecular weight of the protein by 10KDa for each labeled thiol. Blotted membranes were imaged with an Odyssey infrared scanner (LiCor), and the signal was quantified with the scanner’s software.

#### Statistical analyses

Each result presented here was derived from three to six independent experiments. For simple comparisons of two experimental conditions, two-tailed, unpaired t-tests were used. Where variances were not equal, Welch’s correction was used. For comparisons of multiple experimental conditions, one-way or two-way ANOVA was used, and if significant overall, post hoc corrections (Bonferroni or Sidak) for multiple pairwise comparisons were made. P-values less than 0.05 were considered significant.

## Author contributions

F.P. conceived the study, designed and carried out most of computer simulations and analysis. F.P. and R.D.M. designed the *in vitro* and *in vivo* experiments with input from J.T.G. and performed the data analysis. E.F and J.C.A. performed transfections of CaMKII mutants in fly cells. I.J.G. carried coarse grained analysis with input from I.B. and started the molecular dynamics simulations. S.L.C. performed all the histological assays in rat and human tissue. F.P., R.D.M., I.B., J.T.G. and J.C.A. contributed to the drafting of the manuscript. All authors discussed the results, commented and approved the manuscript.

