## Supplementary info for "Disulfide bridge formation prevents CaMKII/Calmodulin interaction in Parkinson’s disease"

### Supplemental Information (DRAFT)

The data that support the findings of this study are available from the corresponding author upon reasonable request.

**Molecular dynamics simulations:**

All simulations in this work were performed using the AMBER15 (**REF-AMBER15**) software package, with the GPU version of PMEMD, and were ran on Nvidia GeForce TITAN-X GPUs. The systems were kept at a temperature of 298 K, using Langevin dynamics with a collision frequency of 2 $ps^{-1}$, and part of the protocol used pressure control, via a weak-coupling Berendsen barostat, with a relaxation time of 2 ps. The SHAKE algorithm was adopted, allowing the use of a 2 fs time step. The protocol followed for minimization and equilibration was: 1) 100 cycles of minimization, using the XMIN method, followed by 5,000 cycles using steepest descent, and another 5,000 steps using conjugate gradient; 2) 1 ns of heating, to 298 K, followed by another ns at constant T and P (1 atm), and finally 20 ns with constant T. The simulation box consisted of a CaMKII dodecamer complex with 8 subunits in close conformation and 4 subunits in open conformation. All-atom simulations were performed in a box with explicit water plus ions to neutralize electrostatic charges. Systems contained more than $1.3\times10^6$ atoms, and all production runs had a time length of over 300ns. (**Total simulation time is ~ 2.6μs**).

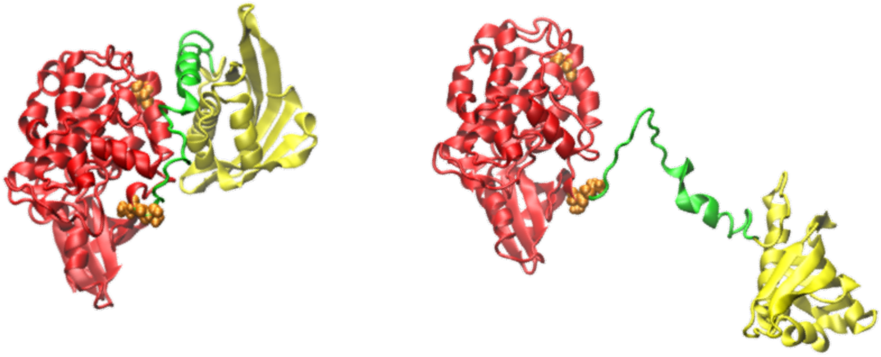

**Figure S1**:
CaMKII monomer. In red the kinase head, in green the linker and in yellow the hub. **Left**, a representation of the closed conformation and **Right,** a conformer of the open conformation obtained in the lab after a combination of free molecular dynamics simulation and umbrella sampling.

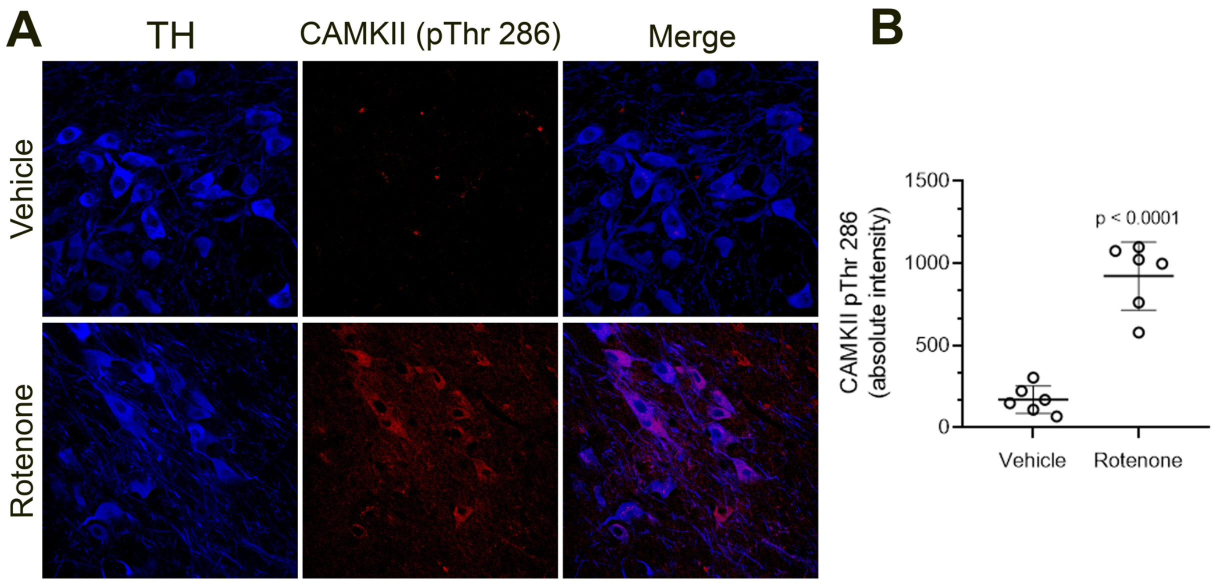


**Figure S2:**
**<u>Thr 286 of CAMKII results hyper phosphorylated in the nigrostriatal neurons of rotenone-treated animals.</u>** Immunohistological analysis of Substantia nigra parsa compacta forma rats treated with rotenone revealed a significant increase of CAMKII phosphoThr 286, when compared to control, suggesting CAMKII hyperactivity of CAMKII under parkinsonian conditions. In each animal, fluorescent signal was measured in 35–50 nigrostriatal neurons per hemisphere. Statistical testing by 2-tailed unpaired t-test with Welch's correction.
